# Deriving stratified effects from joint models investigating Gene-Environment Interactions

**DOI:** 10.1101/693218

**Authors:** Vincent Laville, Timothy Majarian, Paul S. de Vries, Amy R. Bentley, Mary F. Feitosa, Yun J. Sung, DC Rao, Alisa Manning, Hugues Aschard, on behalf of the CHARGE Gene-Lifestyle Interactions Working Group

## Abstract

**Background:** Models including an interaction term and performing a joint test of SNP and/or interaction effect are often used to discover Gene-Environment (GxE) interactions. When the environmental exposure is a binary variable, analyses from exposure-stratified models which consist of estimating genetic effect in unexposed and exposed individuals separately can be of interest. In large-scale consortia focusing on GxE interactions in which only the joint test has been performed, it may be challenging to get summary statistics from both exposure-stratified and marginal (i.e not accounting for interaction) models.

**Results:** In this work, we developed a simple framework to estimate summary statistics in each stratum of a binary exposure and in the marginal model using summary statistics from the “joint” model. We performed simulation studies to assess our estimators’ accuracy and examined potential sources of bias, such as correlation between genotype and exposure and differing phenotypic variances within exposure strata. Results from these simulations highlight the high theoretical accuracy of our estimators and yield insights into the impact of potential sources of bias. We then applied our methods to real data and demonstrate our estimators’ retained accuracy after filtering SNPs by sample size to mitigate potential bias.

**Conclusions:** These analyses demonstrated the accuracy of our method in estimating both stratified and marginal summary statistics from a joint model of gene-environment interaction. In addition to facilitating the interpretation of GxE screenings, this work could be used to guide further functional analyses. We provide a user-friendly Python script to apply this strategy to real datasets. The Python script and documentation are available at https://gitlab.pasteur.fr/statistical-genetics/J2S.

## Background

Gene-Environment (GxE) interactions are of great interest in deciphering biological mechanisms underlying complex human traits and diseases. Several theoretical approaches (1-3) and applications (4-7) have recently been published that identify such GxE interactions. A strategy to detect these interactions applies linear regression models including a GxE interaction term and testing for the hypothesis of null main genetic effect size and GxE interaction effect size, also referred to as the “joint” test (8, 9). Although several interactions have been associated with different traits using this joint test, the main limitation is that of large sample sizes requirements to reach a suitable statistical power (10). The Gene-Lifestyle Interaction Working Group is an international, large-scale, multi-ancestry initiative within the Cohorts for Heart and Aging Research in Genomic Epidemiology (CHARGE) consortium that aims to systematically evaluate genome-wide GxE interactions on cardiovascular disease related traits using genotypic data from up to 610,475 individuals (11). This working group has already unraveled significant GxE interactions using the joint test (12-15). Nevertheless, in the case of binary exposures, alternative approaches can be of interest, notably to identify differential genetic effects between unexposed and exposed individuals. This strategy requires summary statistics computed in each group of individuals separately, which may not always be available in large-scale consortia. Because of logistical challenges, it can be difficult to obtain these summary statistics in such consortia including tens of individual cohorts.

To benefit from these consortia in which only summary statistics in the joint testing framework may be available, we developed a simple tool to infer summary statistics in the groups of unexposed and exposed individuals separately, as well as summary statistics from the regression model without the GxE interaction term. First, we showed that these summary statistics can be efficiently derived from the joint model assuming independence between genotypes and exposure. We then performed a series of simulations to assess the accuracy of these estimations and to examine the impact of different potential sources of bias. Finally, we applied our pipeline to real data from the Gene-Lifestyle Interactions Working group) within the CHARGE Consortium.

### Theoretical derivations

Consider a trait *Y*, a dichotomous exposure *E* and a SNP *G*. A framework to test Gene-Environment interactions is based on the joint model:

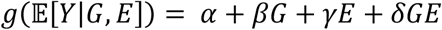

where *g* denotes either the identity function if *Y* is a quantitative trait or the logit function if *Y* is a binary phenotype.

The marginal model in unexposed individuals (E = 0), exposed individuals (E = 1) and all individuals are defined as:

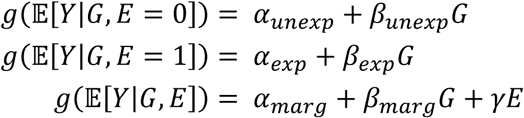

Assuming independence between the genotypes and the exposure (i.e 𝔼[*G*|*E* = 0] = 𝔼[*G*|*E* = 1] = *G*), the joint model can be used to retrieve the marginal genetic effects *β*_*unexp*_ and *β*_*exp*_ in unexposed (*e* = 0) and exposed (*e* = 1) individuals respectively:

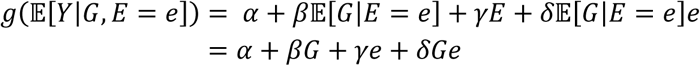

Then setting e to either 0 or 1, marginal effect sizes in each group of individuals can be derived:

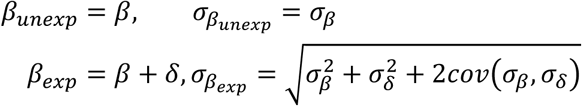

where *σ*_*β*_ and *σ*_*δ*_ denote respectively the standard errors of the genetic effect and interaction effect in the joint model.

Similarly, summary statistics in the marginal model (excluding the interaction term) can be derived from the joint model:

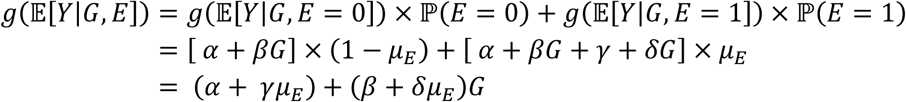

Hence, the marginal genetic effect *β*_*marg*_ and its standard error 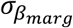 are equal to:

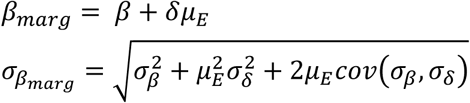

### Implementation

We developed a Python script to derive summary statistics in the marginal model and in each group of individuals separately. As input, the script takes one file with the summary statistics from the joint model, that are genetic and interaction effect sizes, their standard errors, the correlation between the two effect sizes and the sample size for each SNP. This file corresponds to the output of the METAL software to meta-analyze GxE screenings using the joint test (9). In addition to this file, the script also takes two arguments that are the total sample size *N* of the study and the number of exposed individuals *N*_*e*_ included in the study. These two arguments are used to infer the sample sizes 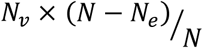 and 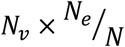 in the group of unexposed and exposed individuals respectively for each SNP, where *N*_*v*_ is the sample size of the SNP. We also implemented a filtering procedure to exclude variants with a low sample size compared to the distribution of the sample sizes: a SNP with a sample size below the 9^th^ decile of the sample size distribution divided by 1.5 are excluded from the analysis. As output, the script generates a single file containing the genetic effect size and its standard error in the group of unexposed individuals, in the group of exposed individuals and in the total sample. The script and a detailed documentation using an example are available at https://gitlab.pasteur.fr/statistical-genetics/J2S.

## Results

### Simulation study

First, we performed a simulation study to assess the accuracy of the estimations obtained from the theoretical results described above. In each of the 1,000 replicates, we simulated 10,000 genotypes of a SNP with a random MAF between 1% and 50% and a binary exposure with a random probability of being exposed ranging from 0.1 to 0.5. Then, we simulated a continuous phenotype *Y* = *β*_*G*_*G* + *β*_*G*_*E* + *β*_*GE*_*G* × *E* + *ε* as a linear combination of the SNP *G*, the exposure *E* and the *G* × *E* interaction term with randomly chosen effect sizes *β*_*G*_, *β*_*E*_ and *β*_*GE*_ and a random noise *ε*∼𝒩(0, *σ*^2^). The effect sizes *β*_*G*_, *β*_*E*_ and *β*_*GE*_ were drawn from a uniform distribution on [0.05; 0.2] with a randomly and equiprobably chosen sign. Note that in this design, genotypes *G* and exposure *E* were drawn independently. Then, we computed the summary statistics from the joint model including the GxE interaction term using individual level data. We also applied linear regressions without the GxE interaction term in each group of individuals (unexposed and exposed) separately and in the pooled sample to compute the summary statistics of the genetic effect in each group of individuals and in the marginal model. Using the estimators derived above, we also inferred these summary statistics in each group and in the marginal model. Comparisons of the empirical and inferred summary statistics showed high accuracy of the estimators, with intraclass correlation coefficient (ICC) between “real” and “estimated” equal to 1 in all scenarios (**Figure 1**).

**Figure 1.**
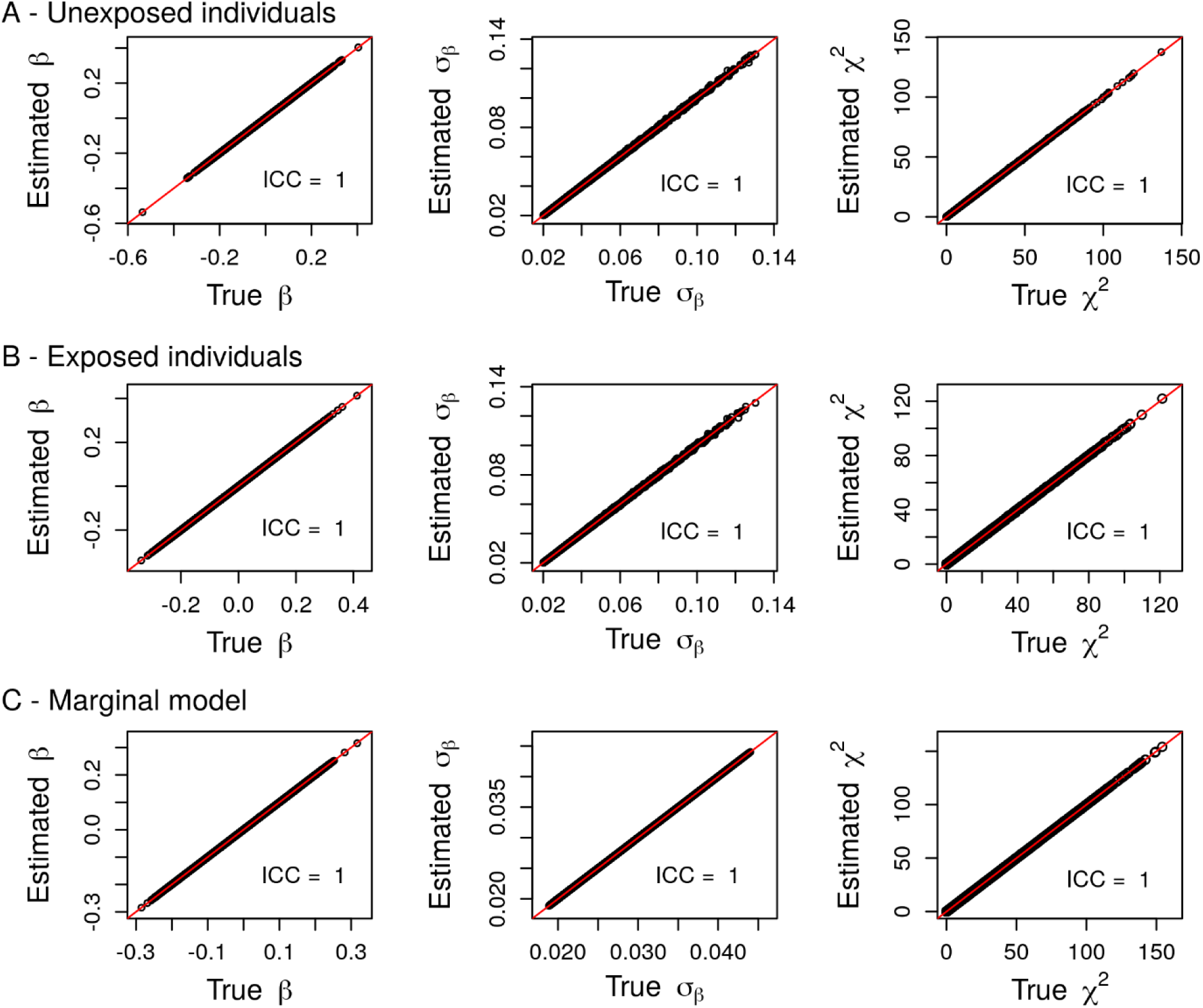
Comparison between summary statistics derived from individual-level data (True) and their estimations (Estimated) in unexposed (A) and exposed (B) individuals and in the marginal model (C) using simulated data in the case of a quantitative phenotype.

We also performed this simulation study for a binary trait. For each of the 10,000 replicates, a quantitative trait *Y*\* = *β*_*G*_*G* + *β*_*G*_*E* + *β*_*GE*_*G* × *E* + *ε* was generated as described above, and we then simulated a binary phenotype *Y* by thresholding *Y*\*. We draw a random number *p* from a uniform distribution ranging in [0.1; 0.4] and set *Y* = 0 if *Y*\* was below the *p*^*th*^ quantile of*Y*\*. As with quantitative traits, the estimator was highly accurate (**Figure S1**).

### Potential bias sources

We performed several complementary simulation studies to assess the contribution of several bias sources.

First, the estimators’ derivation relies on the assumption that genotypes and environment are statistically independent. We performed a simulation study in which correlation existed between genotypes and the environment. We then compared our summary statistics estimated from the joint model to summary statistics derived using individual-level data (**Figure 2, Figure S2**). Relaxing the G-E independence assumption did not impact the estimator’s accuracy when deriving stratified summary. However, estimations in the marginal model were slightly impacted by the correlation between G and E. Indeed, inferred effect sizes are a little biased and effect sizes standard errors are overestimated. Although estimation errors increase with the correlation, the impact on the test statistics remains very limited.

**Figure 2.**
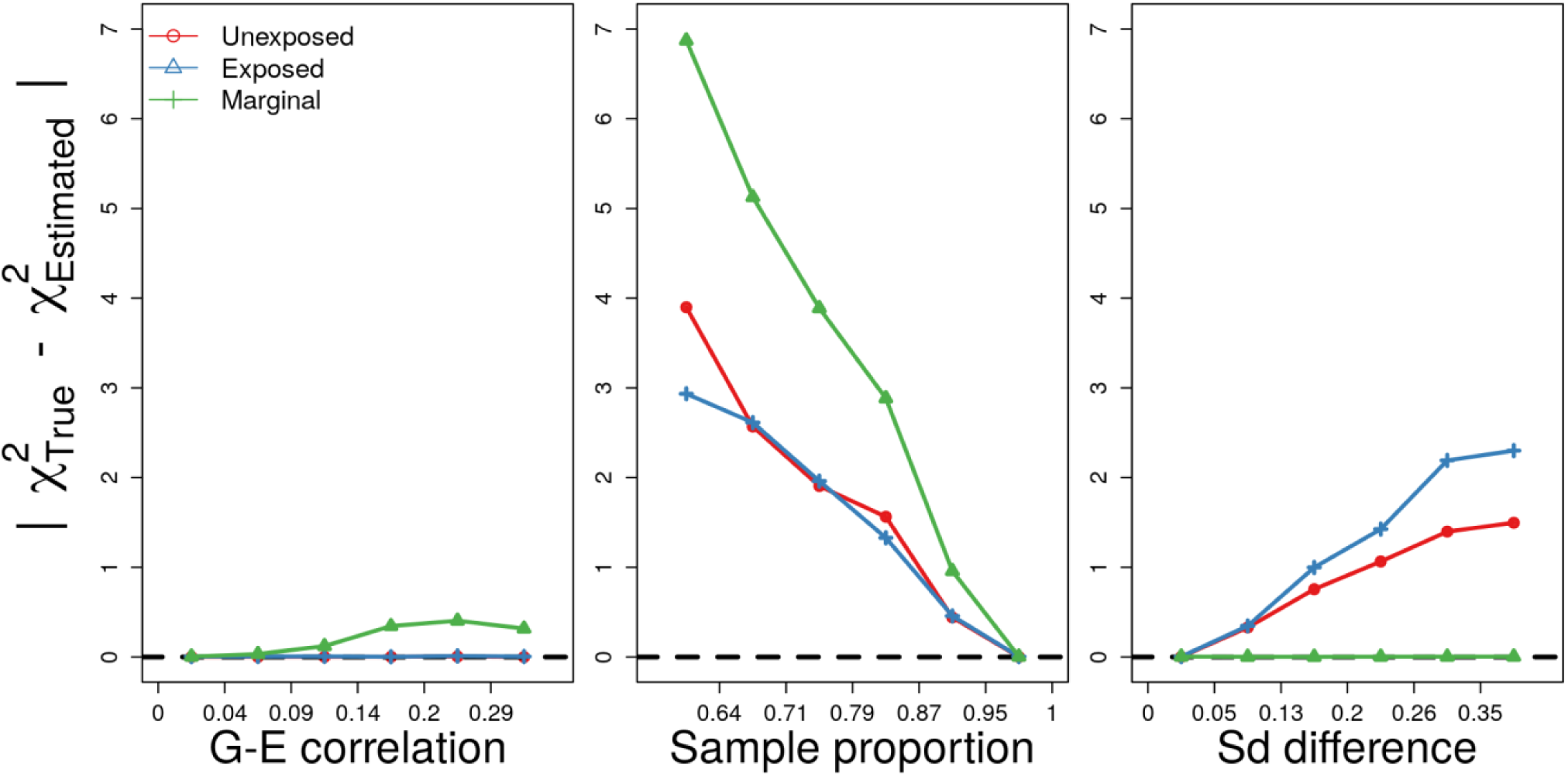
Impact of the different sources of bias on the estimations. The mean of the absolute value of the difference between the test statistics from real data analysis 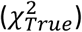 and the test statistics estimated from the summary statistics in the joint model 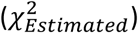 in unexposed individuals only (red), exposed individuals only (blue) and in the marginal model (green) are plotted by quintiles of the G-E correlation coefficient distribution (left), sample proportion distribution (middle) and the distribution of the difference in phenotypic standard deviation between unexposed and exposed individuals (right).

To assess the contribution of low sample size compared to the distribution of sample sizes in the screening, we simulated data for 10,000 individuals and then randomly selected a subset of individuals to estimate the summary statistics using individual-level data. For each SNP, we randomly draw a proportion of individuals to include in the individual-level data model using a uniform distribution ranging between 0.4 and 1. As expected, differences in sample size strongly impact the estimators’ accuracy: greater sample size difference yields less accurate estimations for both the stratified and marginal models (**Figure 2, Figure S3**). The estimation of genetic effect sizes is noisy and their standard errors are systematically underestimated resulting in substantially inflated test statistics.

Finally, bias in our estimations can also occur due to differences in phenotypic variance between unexposed and exposed individuals. To explore this, we simulated a phenotype with exposure-dependent variance by adding statistical noise to the phenotypes of exposed individuals and performed the same simulation study as described above. A different phenotypic variance in the two groups of individuals did not bias estimation of the summary statistics in the marginal model but it clearly biased the estimation of summary statistics in the exposed and unexposed individuals (**Figure 2, Figure S4**). Although this exposure-dependent phenotypic variance did not impact the estimation of the effect sizes, it biased the estimation of the effect size standard error. Standard errors tend to be overestimated in the group in which the phenotypic variance is the largest, leading to deflated test statistics and conversely. Importantly, the larger differences in phenotypic variance yielded larger induced biases.

### Real data application

We assessed the accuracy of our estimations using real data from the Gene-Lifestyle Interaction Working Group of the CHARGE consortium (11). This Working Group recently published genome-wide SNP-by-alcohol interaction screenings (13) using joint tests and focusing on three lipids level: triglycerides (TG), high-density lipoproteins (HDL), and low-density lipoproteins (LDL). Genome-wide screenings for genetic marginal effects were also performed in unexposed and exposed individuals separately and in the whole sample. Here, we used summary statistics from the genome wide SNP by exposure interaction screenings in individuals from European ancestry and derived marginal summary statistics in unexposed and exposed individuals separately, and in the whole sample. We then compared the inferred summary statistics with the empirical summary statistics derived using individual-level data (**Figure 3-5)**. The estimations exhibited high accuracy as demonstrated by the very high ICC between the estimated and true summary statistics (mean ICC = 0.99). Overall, filtering to exclude SNPs with low relative sample size (i.e below the 9^th^ decile of the sample size distribution divided by 1.5) lead to more accurate estimations

**Figure 3.**
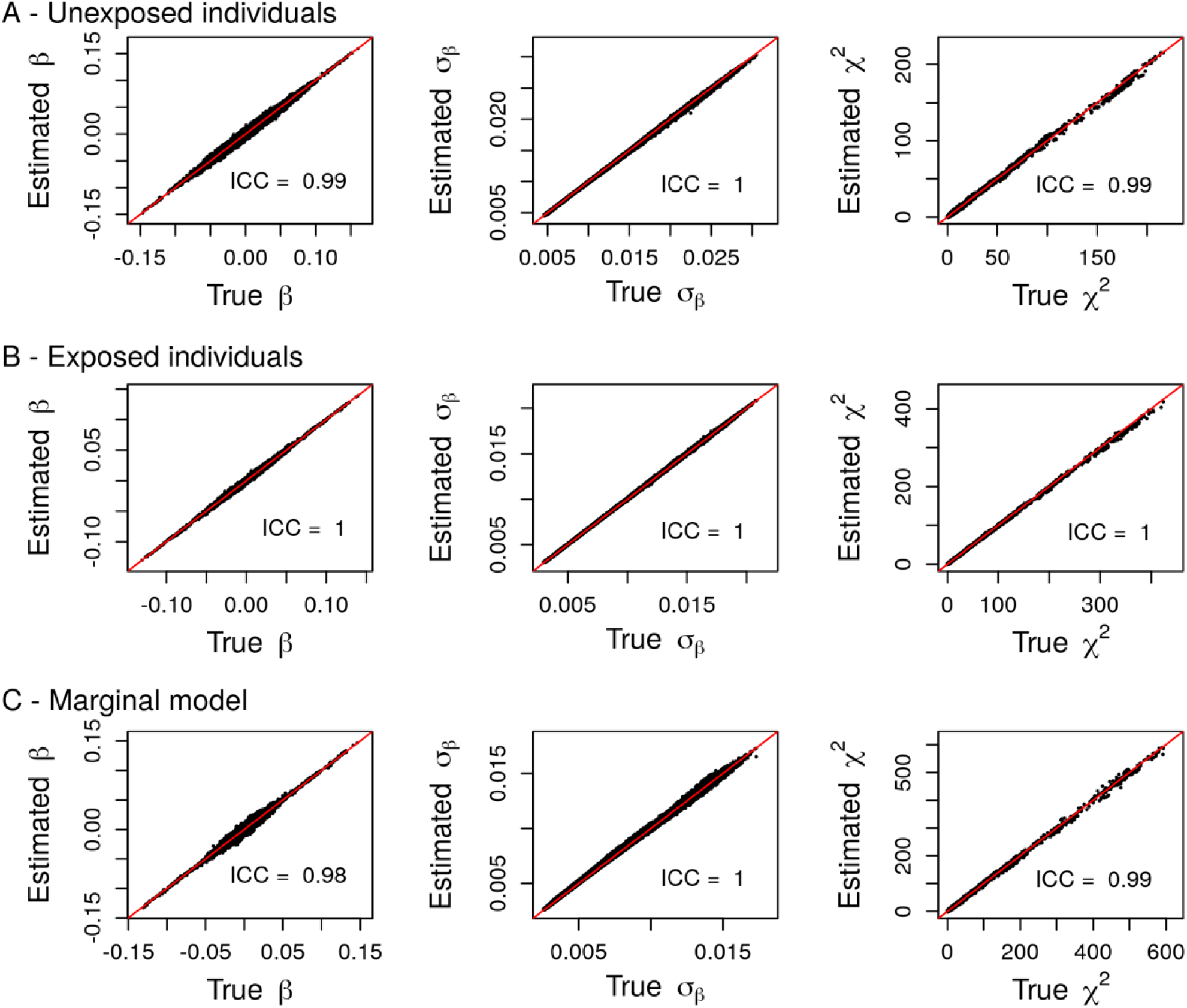
Comparison between summary statistics derived from individual-level data (True) and their estimations (Estimated) in unexposed (A) and exposed (B) individuals and in the marginal model (C) using real data summary statistics from the SNP by alcohol screenings on triglycerides.

**Figure 4.**
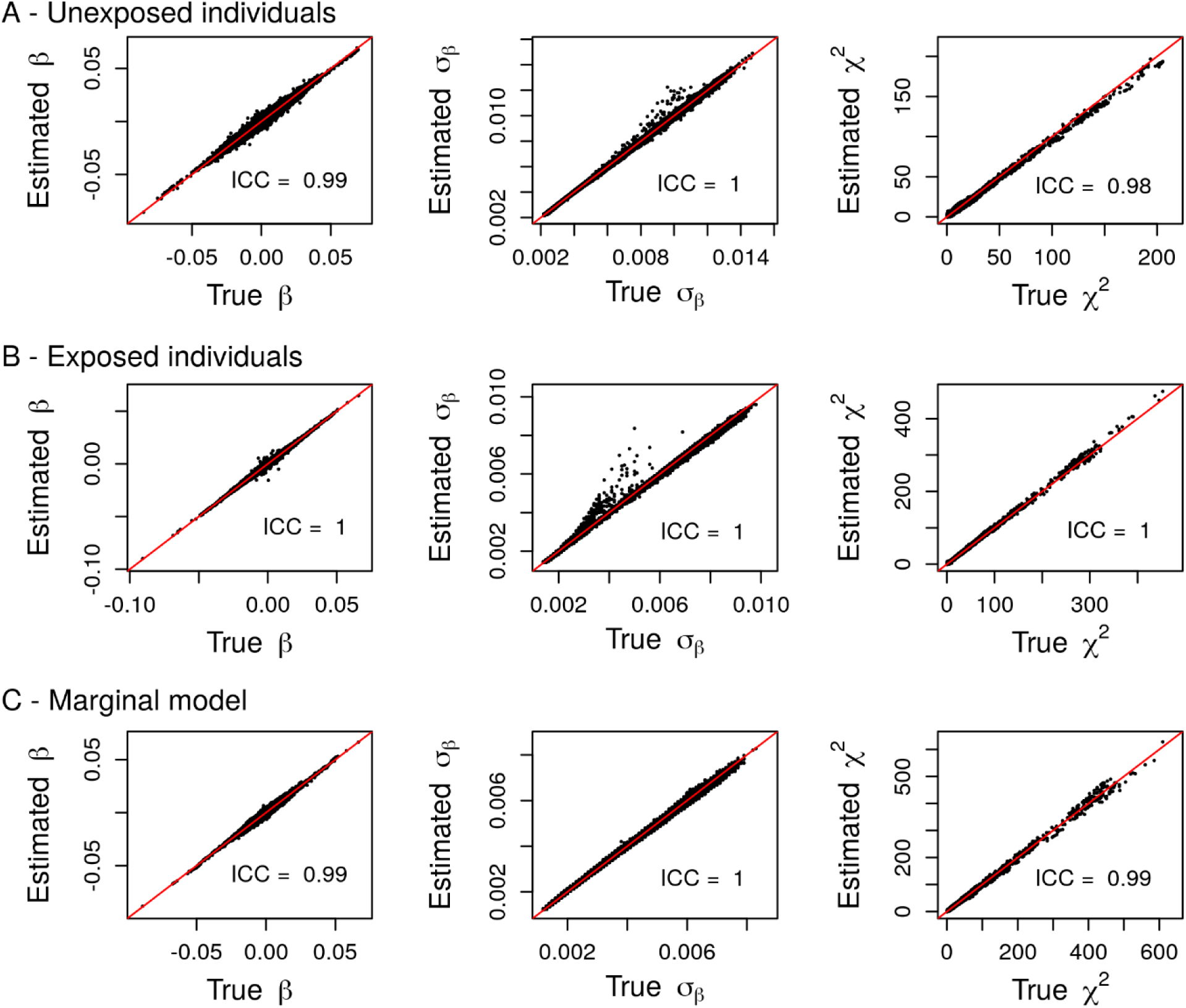
Comparison between summary statistics derived from individual-level data (True) and their estimations (Estimated) in unexposed (A) and exposed (B) individuals and in the marginal model (C) using real data summary statistics from the SNP by alcohol screenings on High Density Lipoproteins.

**Figure 5.**
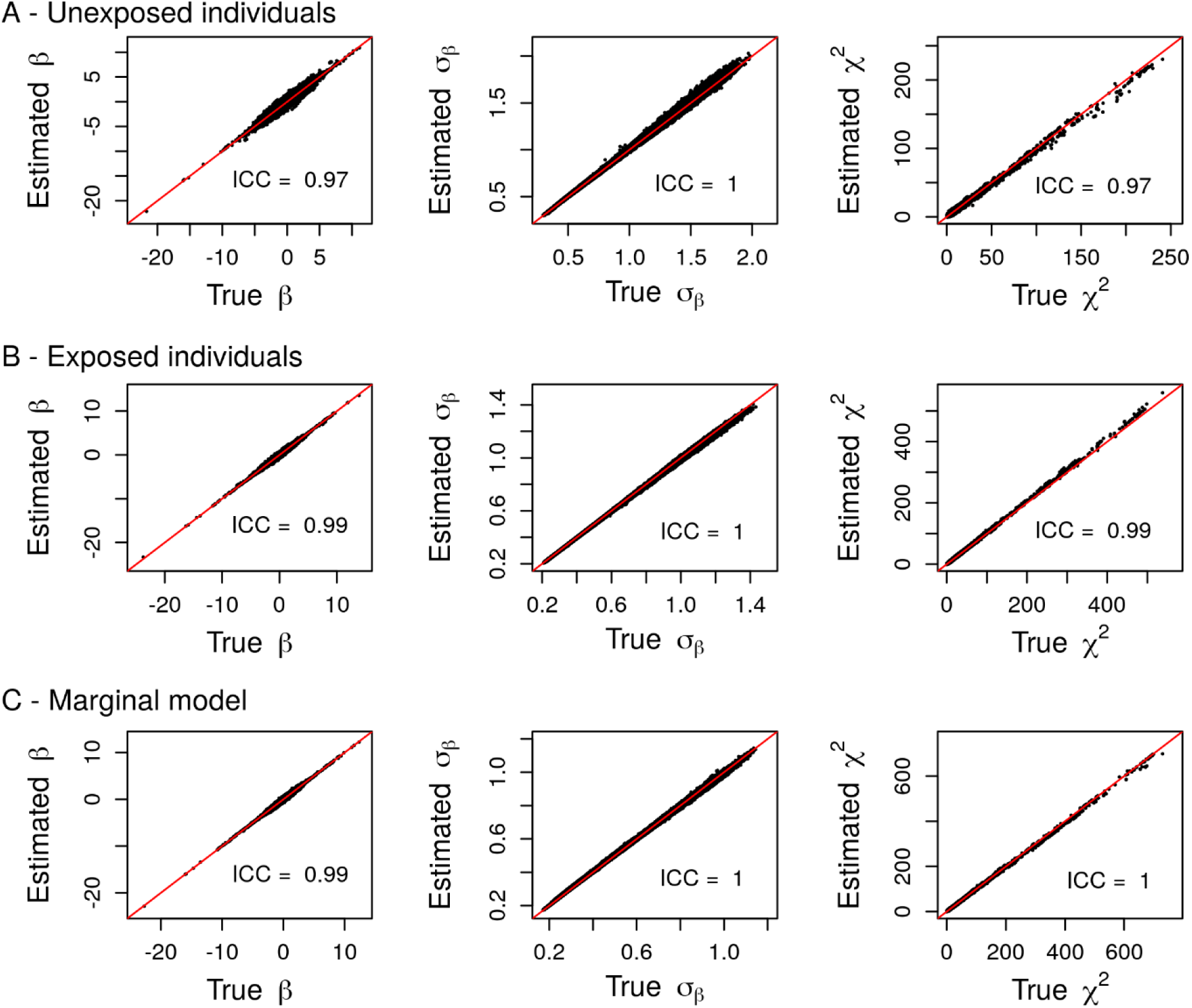
Comparison between summary statistics derived from individual-level data (True) and their estimations (Estimated) in unexposed (A) and exposed (B) individuals and in the marginal model (C) using real data summary statistics from the SNP by alcohol screenings on Low Density Lipoproteins.

## Discussion

In this work, we aimed at inferring marginal genetic effects in exposed and unexposed individuals separately and in the whole sample using summary statistics of the joint test performed in the context of GxE interaction studies. We analytically derived estimators of marginal genetic effects in the different groups of individuals and in the total sample. We validated the method through simulation studies and real data applications which both highlighted the accuracy of our estimations. Notably, this method can be applied without loss of accuracy to quantitative and binary traits.

Expectedly, the most important discrepancies between estimations and real data were observed when the sample size for a SNP differs between the joint model and the stratified models. However, in real data applications in which only the summary statistics from the joint model are available, sample sizes in the stratified models remain unknown. Our method also provides basic estimates of the expected sample size in the groups of exposed and unexposed individuals. Practically, these estimates can be biased for SNPs with a low sample size in the joint test framework because of sampling as the proportion of exposed individuals can be different from its expectation (e.g. if the study includes 30% of exposed individuals, summary statistics for a SNP with a low sample size may have been computed including only 20% of exposed individuals). Consequently, we implemented a procedure to filter out variants with low relative sample size to minimize this potential bias.

Our estimations rely on the independence between genotypes and exposures. Relaxing this assumption leads to small impacts on estimations of the marginal effect size standard deviation in the marginal model. Moreover, in real data applications, this assumption may not hold for only a small proportion of SNPs for which the correlations with the exposure are expected to be low, resulting in little overall impact, as observed when validating our estimators using real data from the Gene-Lifestyle Interaction Working Group.

Finally, we evaluated our estimations in the case of exposure-dependent phenotypic variance. Although our simulations showed clear impacts on the estimations in the stratified models, we noted that the error increased with the magnitude of this difference. In real data applications, such differences in phenotypic variance are expected to be small and should consequently have only a limited impact on the estimations in each exposure stratum. Application to real data sets confirmed this notion as our estimations were highly concordant with real data.

Overall, an advantage of exposure-stratified models is that they allow for a comparison between genetic effects in each group of individuals. This different way of quantifying GxE interactions makes the interpretation more intuitive compared to the joint test by comparing genetic effects between the two groups. In addition, exposure-stratified summary statistics can also be used to apply further analyses such as biological pathways (16) or heritability-based (17-19) analyses. Results from those analyses in each group could then be compared and help better understanding the genetic architecture of the trait. These strategies could also highlight different genetic mechanisms induced by the exposure, opening new path towards public health prevention policies or the identification of potential drug targets.

## Conclusion

In this work, we derived accurate estimations of the marginal genetic effects in unexposed and exposed individuals separately and in the whole sample in the context of genome-wide GxE interaction screenings using the joint test. This method can not only lead to a more intuitive understanding of GxE interactions but also be used to perform additional studies that can guide further functional analyses. We implemented J2S, a Python3 script to easily apply this method, available at https://gitlab.pasteur.fr/statistical-genetics/J2S.

## Availability and requirements

**Project name:** joint2strat

**Project home page:** https://gitlab.pasteur.fr/statistical-genetics/J2S

**Operating systems:** Linux

**Programming language:** Python3

**Other requirements:** None

**License:** MIT

**Any restrictions to use by non-academics:** None

## Abbreviations

GLIWG: Gene-Lifestyle Interaction Working Group
GxE: Gene-Environment
ICC: Intraclass Correlation Coefficient
SNP: Single Nucleotide Polymorphism

## Declarations

### Acknowledgements

The authors acknowledge all the people who involved in the generation and sharing of the data from the CHARGE consortium.

### Authors’contribution

VL and HA designed the study; VL developed the script, VL and TM performed simulation studies and evaluated the accuracy of estimations in real data applications; PSdV, ARB, MFF and YJS generated individual-level data analysis used in the study, DCR, AM and HA supervised the study; VL and HA wrote the manuscript. All authors contributed to the improvement of the manuscript, agreed to be responsible for the accuracy and integrity of this work and provided final approval of the manuscript.

### Ethics approval and consent to participate

Not applicable

### Consent for publication

Not applicable

